# Medial septum activation improves strategy switching once strategies are well learned via bidirectional regulation of dopamine neuron population activity

**DOI:** 10.1101/2022.03.07.483306

**Authors:** D.M. Bortz, C.M. Feistritzer, C.C. Power, A.A. Grace

**Author notes:** Corresponding Author: David M. Bortz, A210 Langley Hall, University of Pittsburgh, Pittsburgh, PA 15260.

## Abstract

Strategy switching is a form of cognitive flexibility that requires inhibiting a previously successful strategy and then switching to a new strategy of a different categorical modality. It is dependent on dopamine (DA) receptor activation and release in ventral striatum and prefrontal cortex, two primary targets of ventral tegmental area (VTA) DA projections. Although the circuitry that underlies strategy switching early in learning has been studied, few studies have examined it after extended discrimination training. This may be important as DA activity and release patterns change across learning, with several studies demonstrating a critical role for substantia nigra pars compacta (SNc) DA activity and release once behaviors are well learned.

Our previous studies demonstrated that medial septum (MS) activation simultaneously increased VTA and decreased SNc DA population activity, as well as improved reversal learning via these actions on DA population activity. We hypothesized that MS activation would improve strategy switching both early in learning and after extended training through its ability to increase VTA DA population activity and decrease SNc DA population activity, respectively. To test this, we activated the MS of male and female rats with designer receptors exclusively activated by designer drugs and measured their performance on an operant-based strategy switching task, following 1, 10, or 15 days of discrimination training. Contrary to our hypothesis, MS activation did not affect strategy switching after 1 day of discrimination training. MS activation improved strategy switching after 10 days of discrimination training, but only in females. MS activation improved strategy switching in both sexes after 15 days of discrimination training. This improvement in strategy switching was attenuated by intra-ventral subiculum bicuculline infusion, which selectively inhibited the MS-mediated decrease in SNc DA population activity, and prevented by infusion of both bicuculline and scopolamine, which inhibited both the MS-mediated decrease in SNc and increase in VTA DA population activity. These data indicate that MS activation improves strategy switching, but only once the original strategy has been sufficiently well learned. They also suggest that the mechanism by which this occurs is likely via the MS’s regulation of DA neuron responsivity, primarily via its ability to down-regulate DA population activity in the SNc.

## Introduction

Cognitive flexibility is the process of adjusting strategies and/or behavior in response to changing environmental or reward contingencies^1^. Strategy switching is one commonly studied form of cognitive flexibility that requires inhibiting a previously successful strategy and then switching to a new strategy that is of a different categorical modality^2–6^. The ability to perform a strategy switch after 1-2 discrimination learning days requires dopamine (DA) receptor activation and release in prefrontal cortex (PFC)^4^ and ventral striatum (VS)^3,6^, two primary targets of ventral tegmental area (VTA) DA projections^7,8^. However, the circuitry that underlies strategy switching after extended discrimination training has not been well studied. This may be important as DA activity and release patterns change across learning. For example, Collins et al (2016) found that DA release in VS predicted learning and performance speed during early learning (1-5 days), but not after extended training (6-7 days)^9^. Furthermore, initiation of an action sequence becomes dependent on the dorsolateral striatum (DLS) once it is well-learned^10^. In fact, the magnitude of phasic DA release from substantia nigra pars compacta (SNc) to the DLS predicts the initiation of a well-learned action^11^ and increases as repetitions increase^12^. This suggests that the DA-related circuitry required to switch strategies after extended training, i.e. once a behavior becomes well learned, may be different from early in learning, and it may include a mechanism for down-regulating SNc to DLS DA activity and/or release.

The medial septum (MS) is a sub region of the basal forebrain that has been most prominently linked to the regulation of theta rhythmicity in the hippocampus^13,14^, and its role in navigation and learning^15–18^. However, two of our previous studies led us to hypothesize that the MS might also play a role in strategy switching. First, MS activation simultaneously increased DA population activity in VTA and decreased it in SNc^19–21^. Because DA population activity regulates the magnitude of DA release^22^, MS activation would have the potential to increase DA release in VTA-projecting regions, such as the VS and PFC, and decrease DA release in SNc-projecting regions, such as the DLS^7,8^. Second, MS activation improved reversal learning, another form of cognitive flexibility, via the abovementioned ability to regulate DA population activity in VTA and SNc^19^. Because of these findings, we hypothesized that MS activation would be able to improve strategy switching both early in learning and after extended training through its ability to increase VTA DA population activity and decrease SNc DA population activity, respectively. We tested this hypothesis by chemogenetically activating the MS and measuring rats’ ability to perform a strategy switch after 1, 10, or 15 discrimination learning days. Next, we replicated^20^ a pharmacological manipulation that allowed us to determine the necessity and individual contribution of the MS’s regulation of VTA and SNc DA population activity for its improvement of strategy switching.

## Materials and methods

### Animals

Experiments were performed using adult female and male Sprague-Dawley rats (250-475g, Envigo, Frederick, MD). Rats were housed in same-sex pairs with ad libitum access to food and water in a temperature- and humidity-controlled room until used. Experimental procedures were approved by the Institutional Animal Care and Use Committee of the University of Pittsburgh according to National Institute of Health Guide for the Care and Use of Laboratory Animals.

### Viral construct and survival surgeries

We used a virus with the Gq-coupled designer receptor exclusively activated by a designer drug (DREADD) attached to the human synapsin promoter and the m-Cherry reporter (DR, AAV_2_ – hSyn – hM3Dq – mCherry; Addgene, Watertown, MA), as well as an empty vector control virus (Con, AAV_2_-hSyn-EGFP, Figure 1/S.1). Prior to any behavioral training, rats were secured to a stereotaxic frame under general anesthesia (isoflurane in oxygen, 5% induction and 2% maintenance) and a burr hole was drilled in the skull. A 5.0 μL Hamilton (Reno, NV) syringe with a 30-gauge needle was lowered at a 5° angle to avoid the sinus and terminated at the more posterior portion of the MS (AP: +0.6mm, ML: +0.55mm, DV: −6.1mm from dura). The syringe was connected to a Micro4 microsyringe pump controller (World Precision Instruments, Sarasota, FL). The viral infusion occurred over 9 minutes (0.1 μL of air + 0.8 μL virus, 0.1 μL/min) with an additional 9 minutes to allow for adequate viral diffusion. Viral spread encompassed the majority of the MS, including on the anterior to posterior plane, as we’ve shown previously^19^. The burr hole was then sealed with bone wax, the incision closed with EZ clips, and rats were temporarily single-housed for recovery. EZ clips were removed and rats were re-paired after a 2-week recovery.

**Figure 1:**
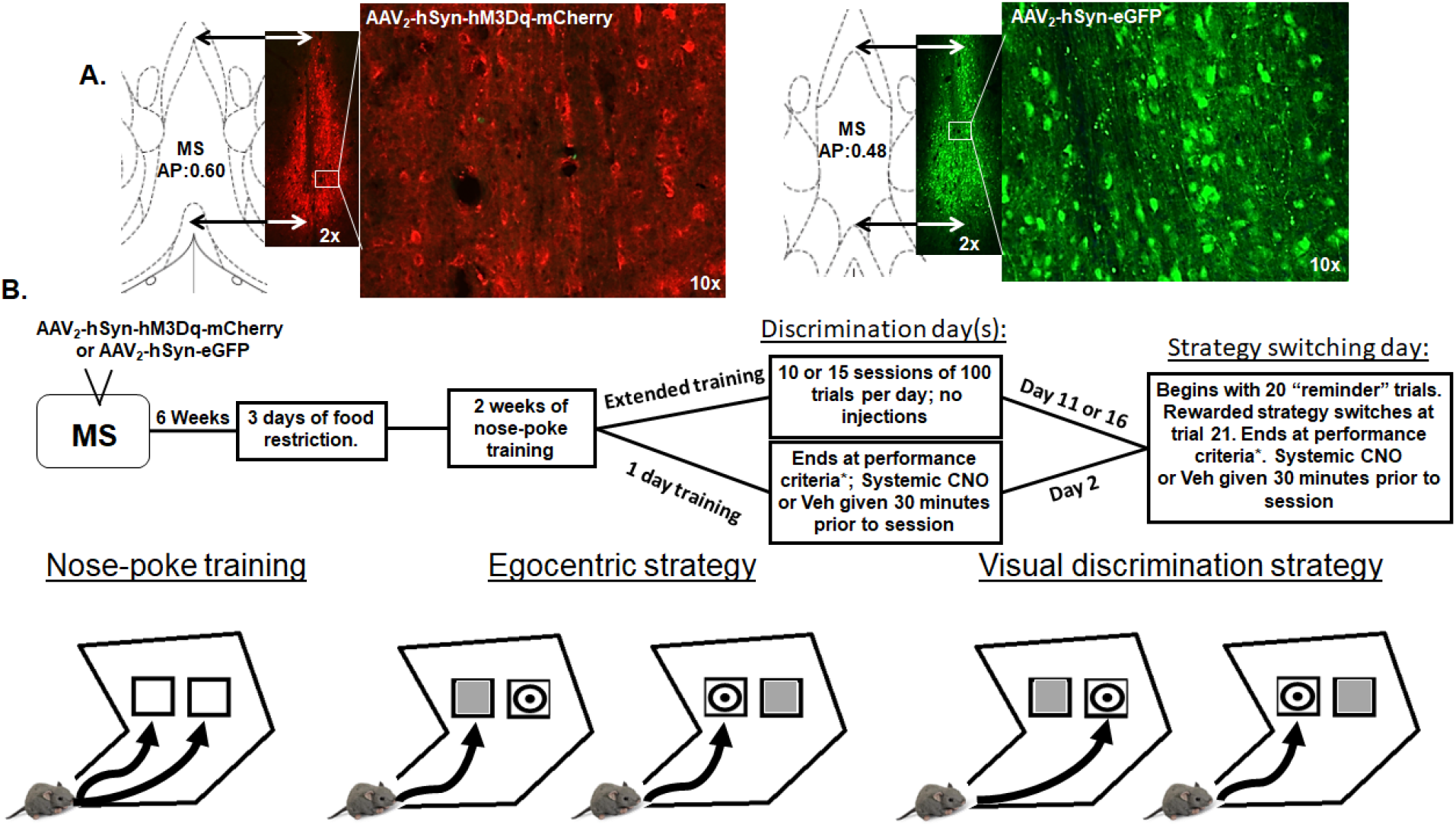
Viral injection, timeline, and testing strategy. **A**. Male and female rats were infused with an hM3Dq-containing (DR, AAV2 – hSyn – hM3Dq – mCherry) or an empty vector control (Con, AAV2-hSyn EGFP) virus into the medial septum (MS, AP: +0.6, ML: ±0.5, DV: −6.1). Both viruses were well-expressed and contained within the MS. Coronal section drawing and associated micrograph illustrates the extent of the fluorescence signal within the MS (2x), and the enlarged micrograph shows expression in neurons within the MS (10x). **B.** Experimental timeline is as follows: Six weeks following the initial viral infusion surgery, all rats were food restricted and trained to nose-poke in response to a visual cue (white square) that appeared in one of two adjacent windows on a touch screen secured to an operant chamber. This “pre-training phase” took approximately two weeks to complete (completion measured by >75% response vs. omission). Rats began the discrimination phase the day following completion of pre-training. All rats were trained on an egocentric discrimination strategy (nose poke only one screen side, regardless of target image) or a visual strategy (nose poke the target image regardless of screen side) and rewarded with a grain pellet. The 1-day training group received a systemic injection of clozapine-N-oxide (CNO; 3 mg/kg) or vehicle (Veh, saline) 30 minutes prior to the discrimination day. *This session continued until the rat reached a performance criterion of 10 consecutive correct trials. The extended training groups performed either 10 or 15 consecutive discrimination training days where no CNO or vehicle injections were given. These lasted for 100 trials per day, regardless of performance. All groups performed a strategy switch test the day after their discrimination day(s) ended (Day 2, 11, or 16). The strategy switching test began with 20 “reminder” trials that rewarded the same strategy that had been learned during the discrimination day(s). Beginning with trial 21, the strategy that was rewarded switched to the other strategy. The test day continued until the rat reached the performance criterion of 10 consecutive correct trials, until a 400-trial max threshold was reached, or 1 hour elapsed. Rats that did not reach criterion in 400 trials or 1 hour continued on subsequent days until criterion was reached. All subsequent test sessions did not include the initial 20 “reminder trials”, but rats did receive an injection of CNO or Veh prior to each additional test day.

### Operant-based strategy switching (Figure 1B)

Six weeks following the initial viral infusion surgery, all rats were food restricted and trained to nose-poke in response to a visual cue (white square). The cue appeared in one of two adjacent windows on a touch screen secured to an operant chamber and appeared evenly on both sides of the screen. Rats performed this stage until they responded reliably to cue presentation on both sides (>75% response vs. omission). Following completion of the pre-training phase, rats learned a complex discrimination where they used either an egocentric strategy (nose poke the left or right screen side regardless of image) or a visual strategy (nose poke the target image regardless of screen side) to earn a grain pellet reward. The 1-day group received a systemic injection of clozapine-N-oxide (CNO, 3 mg/kg) or vehicle (Veh, saline) 30 minutes prior to the discrimination day. This session continued until the rat reached a performance criterion of 10 consecutive correct trials, and there were no differences in learning rates between the two strategy types. The extended training groups performed either 10 or 15 consecutive discrimination training days where CNO or Veh injections were not given. These training days continued for 100 trials per day, regardless of performance. Contrary to the 1-day group, the initial experiments revealed that rats reached a higher degree of performance, at a quicker rate, when learning an egocentric, compared to a visual, discrimination strategy. Thus, all rats in the extended training groups were trained on the egocentric discrimination strategy. All groups performed a strategy switch test the day after their discrimination day(s) ended (Day 2, 11, or 16). The strategy switching test began with 20 “reminder” trials that rewarded the strategy that had been learned during the discrimination day(s). On trial 21, the rewarded strategy switched to the opposite strategy. The test day continued until the rat reached the performance criterion of 10 consecutive correct trials on the new strategy, until a 400-trial max threshold was reached, or 1 hour elapsed. Rats that did not reach criterion in 400 trials or 1 hour continued on subsequent days until criterion was reached. All subsequent test days did not include the initial 20 “reminder trials”, but rats did receive an injection of CNO or Veh prior to each additional test day. In the final experiment, a different group of male and female rats underwent a second surgery to bilaterally implant cannulae into their ventral subiculum (vSub; AP: −6.0mm, ML: + 4.5mm, DL:-8.5mm from skull) just after completion of the pretraining phase. Cannulae were secured with bone screws and dental cement. Rats recovered for 1 week following cannulation surgery and then began the 15-day discrimination training and strategy switch procedure exactly as above. The only difference was that, on test day(s), rats received intra-vSub infusions of vehicle (saline), bicuculline (12.5 ng in 0.5 μL), scopolamine (8 μg in 1.0 μL), or both bicuculline and scopolamine after their systemic injection of CNO or Veh. The test day(s) began at least 30 minutes after the systemic injection and at least 10 minutes after the intra-vSub infusions. Rats that received both scopolamine and bicuculline infusions had at least 5 minutes between infusions. Dependent measures included: total number of trials performed, correct choices, errors, omissions, correct and incorrect choice latency, reward collection latency, premature screen touches (i.e. during the ITI), and error type (never-reinforced vs perseverative). Perseverative errors (e.g. nose-poking the non-target visual cue on the left when the rewarded strategy has switched from “left is correct” to “target is correct”) were compared with never-reinforced errors (e.g. nose-poking the non-target cue on the right when the rewarded strategy has switched from “left is correct” to “target is correct”) to indicate the degree to which the rat was following the previously rewarded strategy vs. exploring in search of the new strategy, respectively.

### Electrophysiological recordings

Following the conclusion of the strategy switching session, DA neuron recordings were performed on all rats using chloral hydrate anesthesia because it has been shown to be most effective in preserving DA neuron activity patterns^23,24^. DA activity in the VTA and SNc was assessed using the 9-electrode track protocol described previously^25,26^ (Figure S.4). Briefly, DA neurons were identified and recorded for 1-3 minutes each using a variety of established criteria, including: low-pitched amplified tone with slow (2-10 Hz) irregular or bursting firing pattern, long duration (2-4ms) biphasic action potential with initial segment-somatodendritic positive phase break, and a temporary inhibition of firing during tail or foot pinch^27–29^. The total number of spontaneously active DA neurons within each animal was counted and then normalized by dividing by the total number of tracks that were examined (active DA neurons per electrode recording track or DA neurons/ track). Three key parameters of identified DA neurons were evaluated: average number of spontaneously active DA neurons per electrode track (calculated for each rat), average DA neuron firing rate, and percent of spikes that occurred in bursts. A burst is defined as beginning with two sequential spikes separated by < 80ms, and ends at > 160ms between spikes. As in the behavioral experiments, recordings began 30 minutes after a systemic injection of Veh or CNO and 10 minutes after intra-vSub infusion of vehicle, scopolamine, bicuculline, or both bicuculline and scopolamine.

### Histology

Rats were euthanized, perfused transcardially with 4% paraformaldahyde for histological analysis of brain sections, and then decapitated. Brains slices were harvested and stained with a mouse mCherry primary (1:8000, Abcam-ab125096) and goat anti-mouse secondary stain (alexa 488, 1:500; Abcam-ab150113) to confirm viral placement and spread. Cannulae placements were histologically confirmed after cresyl violet staining.

### Statistical analyses

In order to properly control for CNO and designer receptor effects, each experiment employed a 4-group design-DREADD virus with CNO injection (DR/CNO), empty vector control virus with CNO (Con/CNO), DREADD virus with vehicle injection (DR/Veh), and control virus with vehicle injection (Con/Veh). For the final intra-vSub infusion experiment, all 4 groups were comprised of rats that received intra-vSub infusions of saline, scopolamine, bicuculline, or both. No intra-vSub infusion had any effect on behavior or DA population activity in the control group rats (Con/CNO, DR/Veh, Con/Veh), as we previously demonstrated^20^. As such, these rats were combined within their respective group as shown in the figures (e.g. Con/Veh/Veh, Con/Veh/Scop, Con/veh/Bicuc, Con/Veh/both = Con/Veh*). All data is reported as mean+SEM and was analyzed with analysis of variance (ANOVA). Post hoc analyses to determine individual group differences were performed using the Tukey’s, Sidak’s, or Fisher’s LSD test (indicated in figure legends).

## Results

### DREADD expression

Viral vectors (DR, AAV_2_ – hSyn – hM3Dq – mCherry or Con, AAV_2_-hSyn-EFP) were highly expressed by cell bodies in the MS, and expression at the infusion site was well contained within the MS (Figure 1A and S.1). Viral transfection also encompassed the vast majority of the anterior-posterior range of the MS, as was shown previously with this same viral infusion protocol^19^. Rats in which viral expression was determined to be lateral or ventral to the MS showed different electrophysiological and behavioral phenotypes than when expression was contained within the MS^19^. Thus, all rats with substantial viral expression outside the MS were excluded from the below analyses.

### MS activation has no effect on discrimination learning or strategy switching after 1 discrimination day

In order to determine the effect of chemogenetic activation of the MS on discrimination learning and strategy switching, a DR or Con virus was infused into the MS of male and female rats (N=14-19 rats/group). Rats performed each day (Figure 1B) 30 minutes after a systemic injection of CNO (3 mg/kg) or Veh (saline). Learning rates were not affected by chemogenetic activation of the MS or sex as all rats had similar trials to reach criterion (Figure 2A, Mean±SEM; Con/Veh: 82.5±13.7, Con/CNO: 80.9±16.9, DR/Veh: 108.3±19.0, DR/CNO: 69.1±14.8; P=0.37) and errors (Con/Veh: 22.6±4.2, Con/CNO: 23.5±5.1, DR/Veh: 32.7±7.1, DR/CNO: 14.7±3.8; P=0.11). Prior to the strategy switch, all rats performed similarly in the 20 “reminder” trials at the beginning of the session. This suggests that rats remembered the rule from the previous day to a similar degree (trials correct out of 20-Con/Veh: 14.9±0.7, Con/CNO: 15.1±0.5, DR/Veh: 14.2±0.7, DR/CNO: 14.9±0.4; P=0.40, not shown). Contrary to our hypothesis, chemogenetic activation of the MS also had no effect on strategy switching (Figure 2B; trials: Con/Veh: 108.2±13.6, Con/CNO: 127.2±25.0, DR/Veh: 136.4±28.5, DR/CNO: 107.2±14.7; P=0.47; errors: Con/Veh: 35.0±4.8, Con/CNO: 40.3±9.3, DR/Veh: 36.2±7.8, DR/CNO: 30.3±4.7; P=0.75). This occurred independently of sex and initial strategy used; thus, the data were combined. These data suggest that chemogenetic activation of the MS is not sufficient to improve strategy switching after a typically used^2–6^ 1-day discrimination-training regimen.

**Figure 2:**
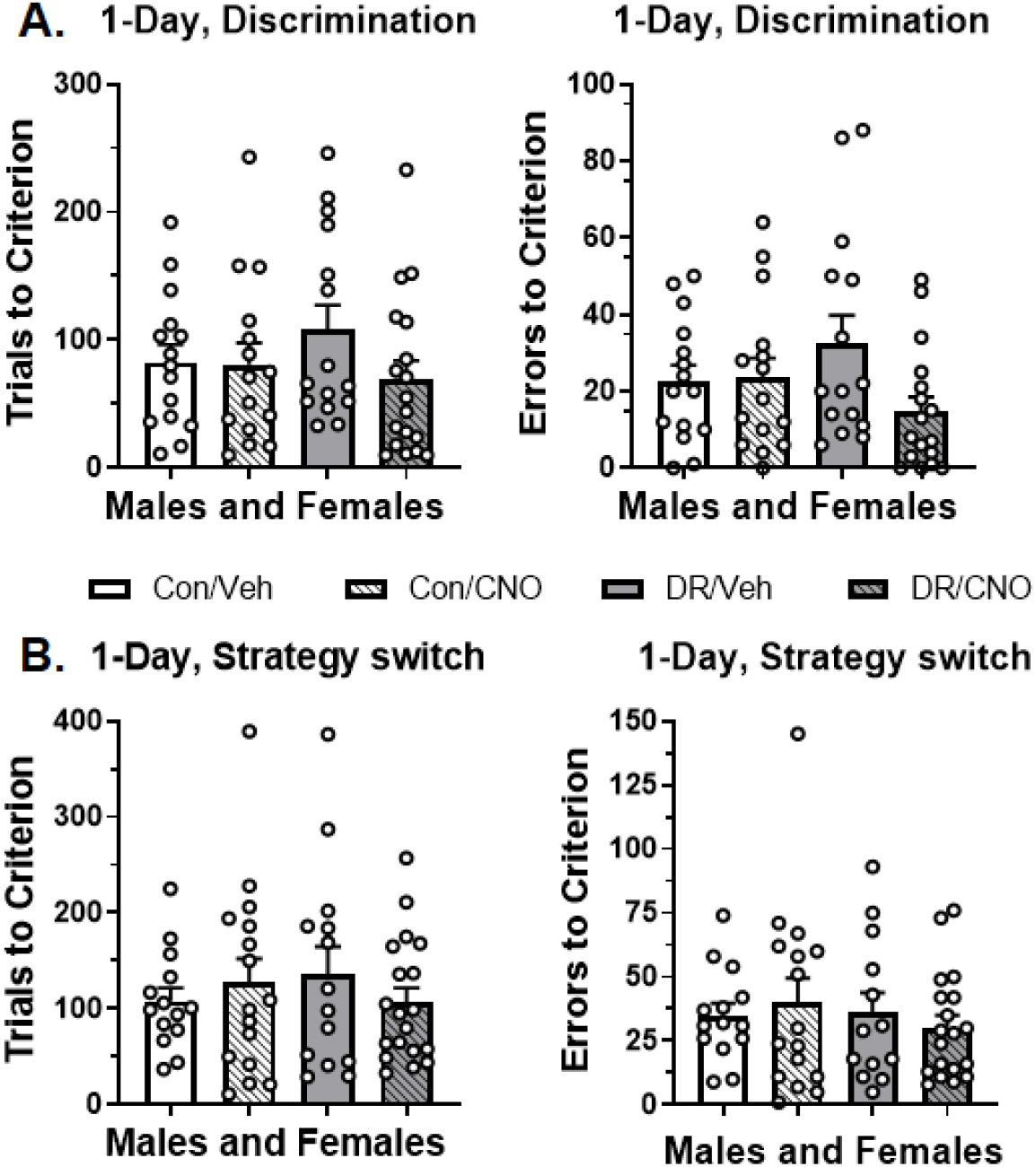
MS activation has no effect on discrimination learning or strategy switching after 1 discrimination day. All experiments in this figure contain both male and female rats (N=14-19 rats/ group), and each open circle represents a data point. **A.** Learning rates were independent of chemogenetic activation of the MS or sex, as all rats had similar trials to reach criterion (*left*, 10 consecutive correct; F_3,59_=1.06, P=0.37) and number of errors committed (*right*, F_3,58_=2.10, P=0.11). **B.** Chemogenetic activation of the MS also had no effect after the strategy switch (after trial 21), both in terms of trials to reach criterion (*left*, 10 consecutive correct; F_3,59_=0.47, P=0.71) and number of errors committed (*right*, F_3,58_=0.40, P=0.75).

### MS activation improves strategy switching after 10 days of discrimination training in female rats

To compare the effect of MS activation on strategy switching after 1 discrimination day vs extended training, a separate set of male and female rats were infused with the DR or Con virus in their MS. Rats were trained on the egocentric strategy for 100 trials per day, for 10 consecutive days, but <u>did not</u> receive CNO or Veh injections during these training days. Analysis of the 10 discrimination days (Figure 3A) revealed that female rats performed significantly better than males on day 3 (P=0.003) and 4 (P=0.013), suggesting that they learned the egocentric discrimination faster (day X sex interaction= P<0.0001). On the 11^th^ day, rats performed the strategy switch test day 30 minutes after a systemic injection of CNO (3 mg/kg) or Veh (N=10 rats per sex, per group). All rats performed similarly in the 20 “reminder” trials at the beginning of the session, suggesting that rats remembered the egocentric strategy to a similar degree (Figure S.2A). Data following the strategy switch on trial 21 showed that MS activation (DR/CNO group) significantly improved female, but not male, rats’ ability to perform the strategy switch, compared to the control groups (Figure 3B). In DR/CNO females, both the number of trials to reach criterion (Con/Veh: 495.4±57.4, Con/CNO: 532.6±49.3, DR/Veh: 532.3±57.2, DR/CNO: 248.3±32.8; F_3,36_=7.46, P=0.0005) and errors (Con/Veh: 178.2±22.8, Con/CNO: 200.3±20.9, DR/Veh: 172.3±22.4, DR/CNO: 82.7±11.6; F_3,36_=6.75, P=0.001) were significantly reduced compared to all three control groups (all p’s < 0.02). Error type analysis revealed that female DR/CNO rats committed fewer perseverative (P=0.002), but not never-reinforced (P=0.13), errors (Figure S.2B). Female DR/CNO rats showed a significant increase in the time to make a correct (P=0.0038), but not incorrect (P=0.11), choice (Figure S.2D). They also showed significantly lower impulsive screen nose-pokes during the inter-trial interval (Figure S.2C, P=0.041). These changes occurred without affecting the latency to collect the pellet reward (Figure S.4E, P=0.24). Although male DR/CNO rats did show a reduction in trials (Con/Veh: 538.1±67.1, Con/CNO: 426.8±57.9, DR/Veh: 422.0±53.7, DR/CNO: 354.0±25.1, F_3,36_=2.05, P=0.12) and errors (Con/Veh: 221.6±33.7, Con/CNO: 174.9±26.0, DR/Veh: 175.7±21.2, DR/CNO: 133.7±9.7; F_3,36_=2.19, P=0.11), neither effect reached statistical significance (Figure 3B). Thus, chemogenetic activation of the MS was sufficient to improve strategy switching after 10 days of discrimination training, but only in females. This improvement was shown as a slowing of decision-making *en route* to switching in fewer trials and committing fewer errors.

**Figure 3:**
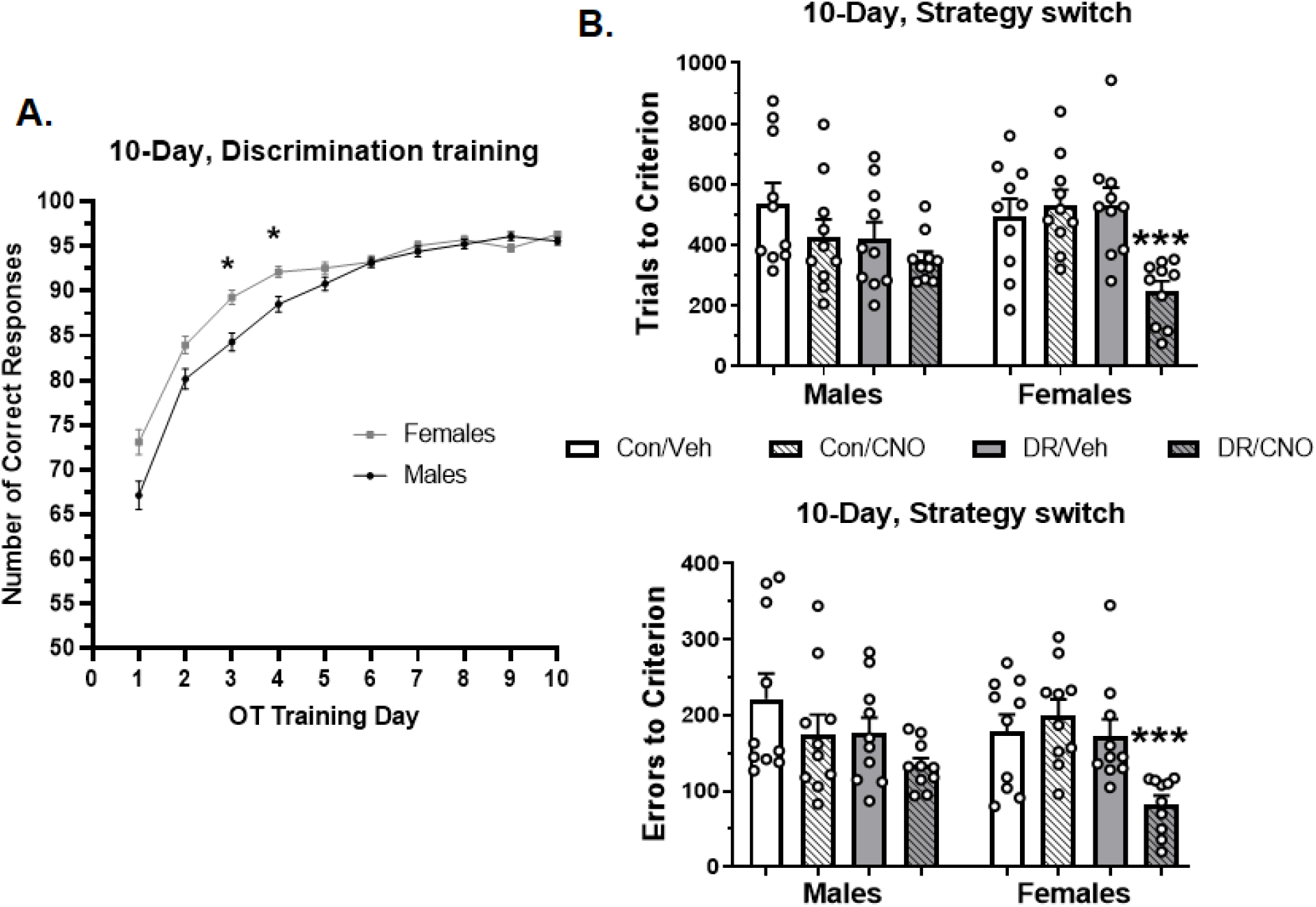
MS activation improves strategy switching after 10 days of training in female rats. Data is split by sex (N=10 rats per group, per sex), and each circle represents a data point. **A.** Female rats learned the egocentric discrimination faster, reaching asymptotic level of performance (defined as > 92% correct) in fewer days than male rats (day X sex interaction: F_9,666_=5.62, P<0.0001). *A post-hoc Sidak’s test revealed this to be the case specifically at day 3 (P=0.003) and 4 (P=0.013). **B.** MS activation (DR/CNO group) in females significantly reduced the number of trials to reach criterion (*top*, 10 consecutive correct, F_3,36_=7.46, P=0.0005) and errors (*bottom*, F_3,36_=6.75, P=0.001) compared to controls. ***A post-hoc Tukey’s test revealed that the female DR/CNO rats were significantly reduced from all 3 of the control groups for both trials (vs. Con/Veh P=0.007, vs. Con/CNO P=0.002, vs. DR/Veh P=0.002) and errors (vs. Con/Veh P=0.009, vs. Con/CNO P=0.001, vs. DR/Veh P=0.016). Although male DR/CNO rats did show a minor reduction in trials and errors, neither effect reached statistical significance.

### MS activation improves strategy switching in both sexes after 15 days of discrimination training

Because female rats learned the egocentric strategy faster in the prior experiment, they spent more days at a high, asymptotic level of performance. I hypothesized that this increased level of training could have contributed to female, but not male, rats’ demonstrated benefit from MS activation. To test this, another set of male and female rats were infused with the DR or Con virus in their MS and trained for 15 consecutive 100-trial days, instead of 10 (Figure S.3A). On the 16^th^ day, rats performed the strategy switch session as above. Rats again performed the vast majority of the 20 “reminder” trials correctly (Figure S.3B). Different from the 10-day discrimination training group, MS activation produced a comparable degree of improvement in strategy switching in both sexes, so they were combined (Figure 4A; N=12-14 rats/group; see Figure S.3C for data separated by sex). MS activation significantly improved strategy switching performance by reducing both trials to reach criterion (Con/Veh: 529.8±61.2, Con/CNO: 616.7±78.4, DR/Veh: 561.3±66.5, DR/CNO: 376.9±32.4; F_3,47_=3.03, P=0.039) and errors (Con/Veh: 207.0±28.9, Con/CNO: 238.9±28.2, DR/Veh: 226.6±32.0, DR/CNO: 128.4±11.5; F_3,47_=3.92, P=0.014; Figure 4A). Similar to the female DR/CNO rats in the prior experiment, DR/CNO rats of both sexes committed fewer perseverative (P=0.010), but not never-reinforced (P=0.93), errors (Figure 4B). Choice latency was significantly increased in male, but not female, rats (Figure S.3E). However, this was observed when the rats made both correct (P=0.005) and incorrect (P=0.007) choices, suggesting a general slowing of decision making, regardless of benefit. Reward collection latency was unaffected by MS activation in either sex (Figure S.3F; P=0.54). Both sexes of the DR/CNO group showed a significant reduction in impulsive screen nose-pokes during the inter-trial interval (P=0.044; Figure S.3D). These data suggest that chemogenetic activation of the MS was sufficient to improve strategy switching similarly in both sexes once discrimination training was extended to 15 days.

**Figure 4:**
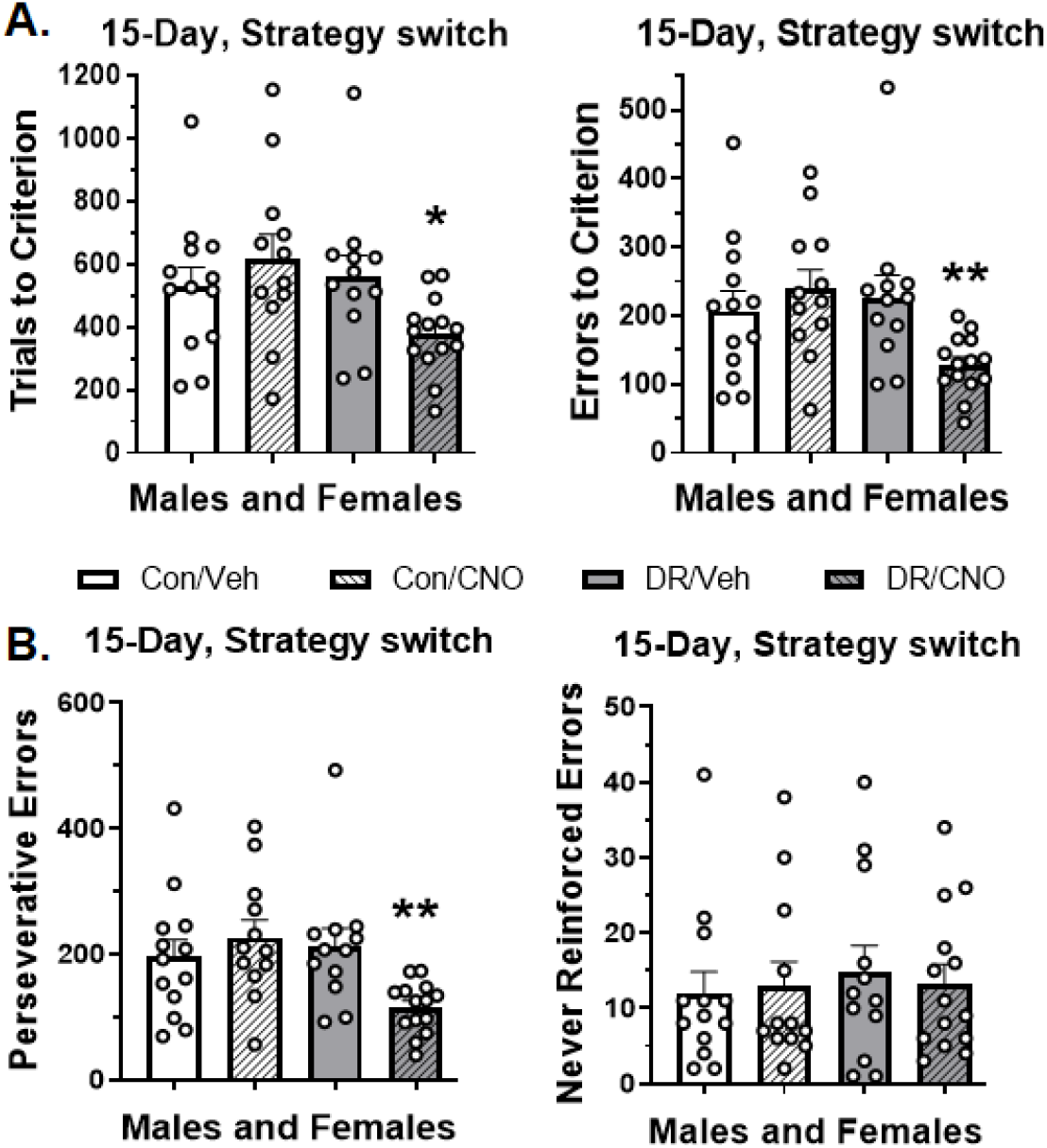
MS activation improves strategy switching in both sexes after 15 days of discrimination training: Performance data of male and female rats (combined) after strategy switch on trial 21 (N=12-14 rats per group). Circles represent each data point. **A.** MS activation (DR/CNO group) significantly improved strategy switching performance in both sexes, as shown by a reduction in both the trials to reach criterion (*left*, 10 consecutive correct; F_3,47_=3.03, P=0.039) and errors (*right*, F_3,47_=3.92, P=0.014). *A Tukey’s test revealed a significant reduction in trials compared to the Con/CNO group (P=0.033), but not the other two (vs. Con/Veh P=0.26 and vs. DR/Veh P=0.14). **A Tukey’s test showed that DR/CNO rat errors were significantly reduced compared to the Con/CNO (P=0.018) and DR/Veh (P=0.043) groups, but not Con/Veh (P=0.13). **B.** DR/CNO rats of both sexes committed fewer perseverative (*left*, F_3,47_=4.26, P=0.010), but not never-reinforced (*right*, F_3,47_=0.14, P=0.93), errors. **A post-hoc Tukey’s tests revealed a significant reduction in perseverative errors compared to the Con/CNO (P=0.012) and DR/Veh (P=0.036) groups, but only a trend compared to the Con.Veh group (P=0.099).

### MS activation affects DA activity similarly in male and female rats and is blocked by pharmacological manipulation of the ventral subiculum

We hypothesized that MS activation was improving strategy switching via its known ability to regulate DA activity in males^19–21^, similar to what we previously showed with reversal learning^19^. To first determine if MS activation affects DA population activity in female rats to the same degree as male rats, we recorded DA activity in females that had finished the strategy switching paradigm (N=5-7 rats/group; see Figure S.4). Female control rats showed similar numbers of spontaneously active DA neurons in the SNc as what has been previously reported^19,20^ in males (Figure 5A; DA neurons/ track-Con/Veh: 1.8 ±0.1, Con/CNO: 1.7±0.1, DR/Veh: 1.7±0.1). Chemogenetic activation of the MS significantly reduced the number of spontaneously active DA neurons in the SNc (DR/CNO: 0.9±0.2; F_3,20_=13.5, P<0.0001), and this, again, was similar to the reduction reported in males^19,20^. This was also true when measuring in the VTA^19,20,30^ for both female control rats (Figure 5B; Con/Veh: 0.8±0.1, Con/CNO: 0.6±0.2, DR/Veh: 1.0±0.1) and after chemogentic activation of the MS (DR/CNO: 1.6±0.2; F_3,21_=8.05, P=0.0009). MS activation did not significantly alter DA neuron firing rate or burst activity (not shown), as has been shown previously^19–21^.

**Figure 5:**
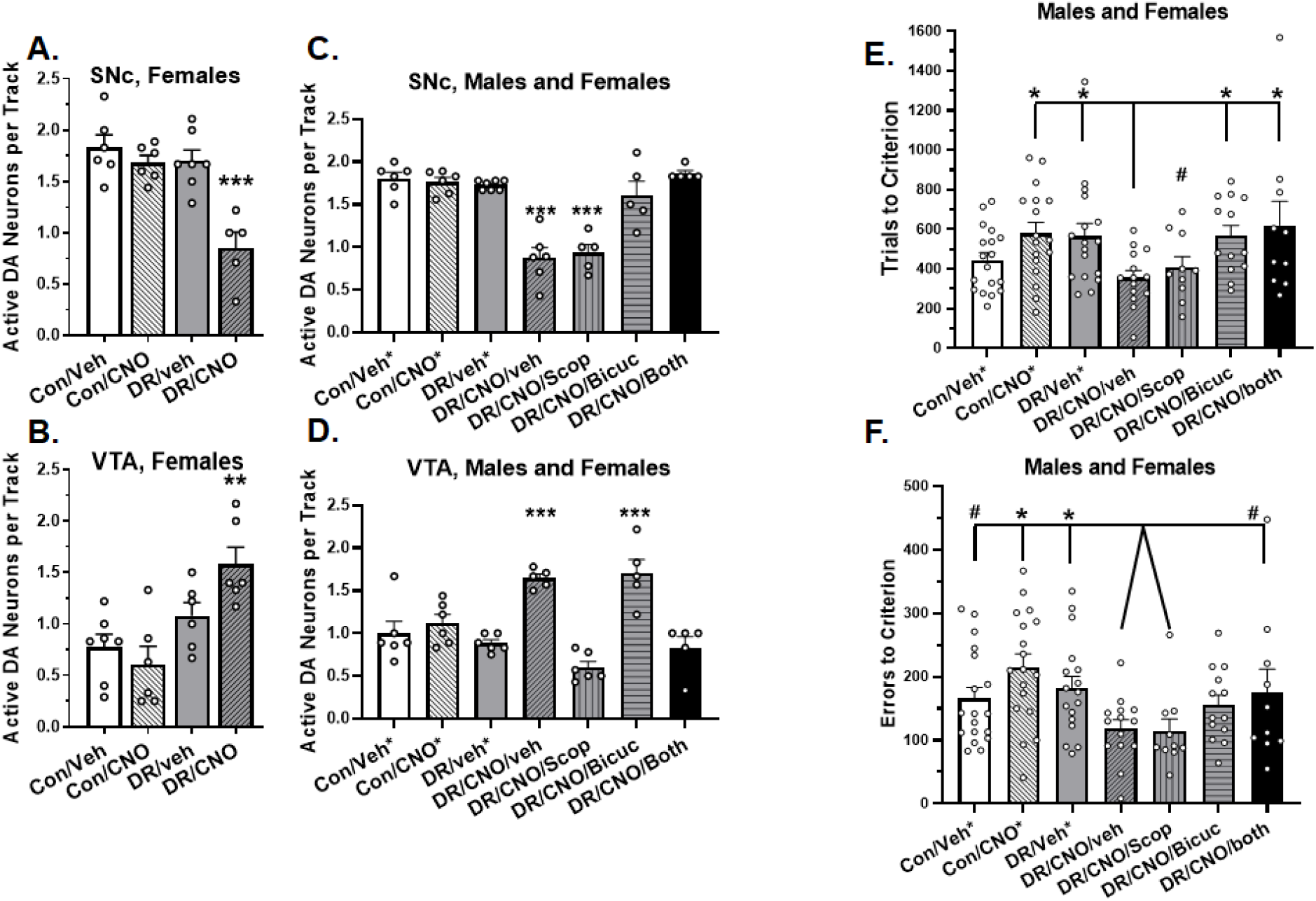
Improvement in strategy switching is primarily mediated via the MS’s ability to decrease DA population activity in SNc. Circles represent each data point. **A.** Chemogenetic activation of the MS (DR/CNO group) significantly reduced DA neuron population activity in the SNc in female rats (N=5-7 rats/ group; F_3,20_=13.5, P<0.0001). ***A Tukey’s test revealed that DA population activity was reduced compared to all three control groups (vs. Con/Veh P<0.0001, vs. Con/CNO P=0.0005, vs. DR/Veh P=0.0002). **B.** Chemogenetic activation of the MS (DR/CNO group) significantly increased DA neuron population activity in the VTA in female rats (N=5-7 rats/ group; F_3,21_=8.05, P=0.0009). **A Tukey’s test revealed that DA population activity was increased compared to Con/Veh (P=0.0043) and Con/CNO (P=0.001), but not DR/Veh (P=0.12). **C.** Intra-vSub infusions had no effect on baseline DA population activity when given to control rats and were combined (e.g. Con/Veh/Veh, Con/Veh/Scop, Con/veh/Bicuc, Con/Veh/both = Con/Veh*, etc.; N=5-7 rats/ group). The reduction in SNc DA population activity after MS activation was prevented by intra-vSub infusion of bicuculline (Bicuc) and both bicuculline and scopolamine (Both), but not by scopolamine (Scop). ***This effect was significant (F_6,33_=22.6, P<0.0001), with a Tukey’s test showing that the DR/CNO/Veh and DR/CNO/Scop groups were significantly decreased from all other groups (vs. Con/Veh* P’s<0.0001, vs. Con/CNO* P’s<0.0001, vs. DR/Veh* P’s<0.0001, vs. DR/CNO/Bicuc P<0.0001 and P=0.0003, vs. DR/CNO/Both P’s<0.0001). **D.** The increase in VTA DA population activity after MS activation was prevented by intra-vSub infusion of scopolamine (Scop) and both bicuculline and scopolamine (Both), but not by bicuculline (Bicuc). ***This effect was significant (F_6,32_=14.5, P<0.0001), with a Tukey’s test showing that the DR/CNO/Veh and DR/CNO/Bicuc groups were significantly elevated from all other groups (vs. Con/Veh* P=0.003 and 0.001, vs. Con/CNO* P=0.023 and 0.009, vs. DR/Veh* P=0.0003 and 0.0001, vs. DR/CNO/Scop P’s<0.0001, vs. DR/CNO/Both P=0.0003 and <0.0001, respectively). **E.** The reduction in trials to reach criterion (10 consecutive correct) after MS activation was inhibited by intra-vSub infusion of bicuculline (Bicuc), prevented by infusion of both bicuculline and scopolamine (Both), but not affected by infusion of scopolamine (N=10-18 rats/ group; Scop; overall F_6,92_=2.63, P=0.021). *A Fisher’s LSD test showed that the DR/CNO/Veh group was reduced compared to the Con/CNO* (P=0.007), DR/Veh* (P=0.012), DR/CNO/Bicuc (P=0017), and DR/CNO/Both (P=0.006) groups. #DR/CNO/Scop rats remained similarly reduced as the DR/CNO/Veh group, showing a trend toward a reduction compared to Con/CNO* (P=0.062), DR/Veh* (P=0.087), and DR/CNO/Bicuc (P=0.0998) groups, and a reduction compared to the DR/CNO/Both group (P=0.040). **F.** The reduction in errors after MS activation was prevented by infusion of both bicuculline and scopolamine (Both; F_6,93_=3.07, P=0.009). Infusion of bicuculline showed a small inhibition of the reduction in errors, but this did not reach significance (vs. DR/CNO/Veh P=0.22). Infusion of scopolamine had no effect on the reduction in errors. A Fisher’s LSD test showed that the DR/CNO/Veh and DR/CNO/Scop groups were *significantly reduced compared to the Con/CNO* (P=0.0006 and 0.0013, respectively) and DR/Veh* (P=0.023 and 0.029, respectively) groups and #showed a trend towards a significant reduction compared to the Con/Veh* (P=0.087 and 0.094, respectively) and DR/CNO/Both (P=0.079 and 0.082, respectively) groups.

We previously showed that the MS regulates DA population activity via a direct projection to the ventral subiculum (vSub)^20^, which then follows a known pathway^31–34^ to the VTA and SNc^20^. Furthermore, we showed that infusion of scopolamine into the vSub selectively inhibited the increase in DA population activity in the VTA after pharmacological MS activation, but did not affect the decrease in SNc^20^. Infusion of bicuculline into the vSub selectively prevented the decrease in SNc, but did not affect the increase in VTA^20^. To confirm these findings chemogenetically, we used male and female rats that had been infused previously with the DR or Con virus in the MS and had completed behavioral experiments. Anesthetized rats were given a systemic injection of CNO or Veh and an intra-vSub infusion of scopolamine (Scop; 8μg in 1μL), bicuculline (Bicuc; 12.5ng in 0.5μL), or both 30 and 10 minutes prior to recording, respectively (N=5-7 rats/group). Similar to what had been shown previously^20^, intra-vSub infusions of bicuculline, scopolamine, or both had no effect on baseline DA population activity when given to control rats. Because of this, they were combined within each control group (e.g. Con/Veh/Veh, Con/Veh/Scop, Con/veh/Bicuc, Con/Veh/both = Con/Veh*). DA population activity in the SNc was similar in male and female rats to what has been shown previously (Figure 5C; Con/Veh*: 1.8 ±0.1, Con/CNO*: 1.8±0.1, DR/Veh*: 1.7±0.02), and was significantly decreased following chemogenetic activation of the MS (DR/CNO/Veh: 0.9±0.1). Intra-vSub infusion of bicuculline or both drugs prevented the decrease in SNc DA population activity (DR/CNO/Bicuc: 1.6±0.2, DR/CNO/both: 1.9±0.03; F_6,33_=22.6, P<0.0001), but scopolamine did not (DR/CNO/Scop: 0.9±0.1). DA population activity in the VTA in male and female rats was also similar to what was shown previously (Figure 5D; Con/Veh*: 1.0±0.1, Con/CNO*: 1.1±0.1, DR/Veh*: 0.9±0.04), and was significantly elevated following chemogenetic activation of the MS (DR/CNO/Veh: 1.6±0.1). Intra-vSub infusion of scopolamine or both drugs prevented the increase in VTA DA population activity (DR/CNO/Scop: 0.6±0.1, DR/CNO/both: 0.8±0.1; F_6,32_=14.5, P<0.0001), but bicuculline did not (DR/CNO/Bicuc: 1.7±0.2). No manipulation significantly affected DA neuron firing rate or burst activity (not shown), as has been shown previously^19–21^. These data confirmed that MS activation affected DA population activity comparably in male and female rats. They also demonstrated that pharmacological manipulation of the vSub was suitable to determine the necessity and individual contribution of the MS’s regulation of VTA and SNc DA population activity for its improvement of strategy switching.

### MS activation-induced improvement in strategy switching is primarily mediated via the ability to decrease DA population activity in SNc

To determine the degree to which the MS’s regulation of DA population activity plays a role in its ability to affect strategy switching, a final set of male and female rats were infused with the DR or Con virus in their MS and bilaterally implanted with cannulae in vSub. Rats followed the 15-day discrimination training procedure outlined above (Figure S.5A). On the 16^th^ day, rats were given a systemic injection of CNO or Veh, as well as bilateral vSub infusions of vehicle (Veh, saline), scopolamine (Scop, 8μg in 1μL), Bicuculline (Bicuc, 12.5ng in 0.5μL), or both 30 and 10 minutes before the test, respectively. Again, vSub infusions had no effect on control performance and were combined as above (N=10-18 rats/group). All rats performed the majority of the 20 “reminder” trials correctly (Figure S.5B). DR/CNO/Scop and DR/CNO/both groups showed a slight, but significant (P=0.005), reduction in number of correct trials out of 20, although this did not affect their performance during the strategy switch. After the strategy switch, MS activation (DR/CNO/Veh) again reduced both trials to reach criterion (Figure 5E; Con/Veh*: 444.4±38.0, Con/CNO*: 579.2±64.5, DR/Veh*: 564.9±64.5, DR/CNO/Veh: 355.1±37.7; DR/CNO/Veh vs. Con/Veh* P=0.27, vs. Con/CNO* P=0.007, vs. DR/Veh* P=0.012) and errors (Figure 5F; Con/Veh*: 165.8.0±17.6, Con/CNO*: 215.3±20.9, DR/Veh*: 182.6±18.4, DR/CNO/Veh: 118.3±13.9; DR/CNO/Veh vs. Con/Veh* P=0.087, vs. Con/CNO* P=0.0006, vs. DR/Veh* P=0.023). Intra-vSub infusion of scopolamine did not affect the reduction in either trials-to-criterion (DR/CNO/Scop: 408.6±53.7; vs. DR/CNO/Veh P=0.57) or errors (DR/CNO/Scop: 114.3±19.3; vs. DR/CNO/Veh P=0.90), but remained similarly reduced compared to the control groups. Intra-vSub infusion of bicuculline significantly reduced the improvement in strategy switching. DR/CNO/Bicuc rats required as many trials to reach criterion as the control rats (567.1±52.9; vs. Con/Veh* P=0.14, vs. Con/CNO* P=0.89, vs. DR/Veh* P=0.98), and significantly more than DR/CNO/Veh rats (P=0.017). Interestingly, DR/CNO/Bicuc rats did not commit significantly more errors compared to DR/CNO/Veh rats (155.2±16.1; P=0.22). Intra-vSub infusion of both bicuculline and scopolamine prevented the improvement in strategy switching mediated by MS activation. DR/CNO/both rats also had comparable trials-to-criterion as the control groups (619.9±122.0; vs. Con/Veh* P=0.053, vs. Con/CNO* P=0.65, vs. DR/Veh* P=0.54), which were significantly increased compared to DR/CNO/Veh rats (P=0.006). However, DR/CNO/both rats also committed a similar number of errors compared to the control groups (174.9±37.0; vs. Con/Veh* P=0.76, vs. Con/CNO* P=0.19, vs. DR/Veh* P=0.80), which trended towards a significant increase compared to both the DR/CNO/Veh (P=0.079) and DR/CNO/Scop (P=0.082) groups. MS activation (DR/CNO/Veh) improved performance by reducing perseverative (vs. Con/Veh* P=0.048, vs. Con/CNO* P=0.0003, vs. DR/Veh* P=0.019), but not never-reinforced (overall P=0.18), errors (Figure S.5C). This effect was blocked only by infusion of both bicuculline and scopolamine (P=0.085), which did not affect never-reinforced errors. Infusion of scopolamine, bicuculline, or both in the DR/CNO, but not control, rats led to a slight, but statistically significant, increase in choice and reward collection times compared to the other 4 groups (Figure S.5E, p’s<0.05). These data demonstrate that the MS’s ability to reduce SNc DA population activity is likely the primary mechanism by which it improves strategy switching.

## Discussion

Chemogenetic activation of the MS had no effect on strategy switching after 1 day of discrimination training. MS activation improved strategy switching after 10 days of discrimination training, but only in female rats. Female rats performed the strategy switch faster (fewer trials) and with fewer mistakes, compared to controls. MS activation improved strategy switching in both sexes after 15 days of training. This improvement in strategy switching was attenuated by intra-vSub bicuculline infusion, which selectively inhibited the MS-mediated decrease in SNc DA population activity, and prevented by infusion of both bicuculline and scopolamine, which inhibited both the MS-mediated decrease in SNc and increase in VTA DA population activity. These data indicate that MS activation improves strategy switching, but only once the original strategy has been sufficiently well learned. They also suggest that the mechanism by which this occurs is likely via the MS’s regulation of DA neuron responsivity, primarily via its ability to down-regulate DA population activity in the SNc.

### Are the sex differences in strategy switching improvement a result of differences in MS function?

Chemogenetic activation of the MS significantly improved strategy switching performance in female rats after 10 days of discrimination training, but not in males. It is possible that this finding reflects a sex difference in MS function; however, few studies have compared MS functionality in male and female rodents, so there is little support for this claim. Our data indicate that MS activation had a comparable effect on male and female DA population activity, even in terms of magnitude of the effect. We also showed that a manipulation that prevented the decrease in SNc and increase in VTA DA population activity also prevented the MS activation-induced improvement in strategy switching in both sexes, indicating that the MS’s regulation of DA population activity is the probable mechanism by which the MS improved strategy switching. Thus, we found no mechanistic evidence that supports distinct MS functionality by sex. Alternatively, a well-performed study by Chen et al (2020) demonstrated that female rats tended to use an egocentric strategy when learning to discriminate between two novel stimuli^35^. Male rats tended to erratically shift strategies based on feedback from the previous trial^35^. This led to female rats learning discriminations faster, as was also seen in the 10-day group. Therefore, a likely explanation for female rats’ faster discrimination learning in the 10-day group is their natural inclination towards the egocentric strategy, which consequently led to a higher degree of training. Thus, we propose that the initial strategy was sufficiently well learned for MS activation to be effective after 10 days of training in the females, but, because of their slower rate of discrimination learning, this was not the case in males. Interestingly, extending discrimination training to 15 days showed a significant benefit of MS activation for strategy switching in both sexes, which supports this notion.

### Proposed mechanism for the MS-mediated improvement in strategy switching

MS activation-induced improvement in strategy switching was attenuated following intra-vSub infusion of bicuculline and prevented following infusion of both bicuculline and scopolamine. It was not affected by infusion of scopolamine alone. Bicuculline selectively prevents the MS activation-induced decrease in SNc DA population activity, while scopolamine selectively prevents the increase in VTA DA population activity. This suggests that the primary mechanism by which the MS improves strategy switching is via its ability to downregulate DA population activity in the SNc. This leaves two questions to be discussed:

#### How does the MS downregulating DA population activity in SNc affect strategy switching?

As rewarded behaviors are repeated over time, and learning progresses, motor action plans become stereotyped^36–38^. DA release that is initially seen in VS and is predictive of learning rate^8,9^, begins to dissipate^12^. This coincides with the emergence of a DA signal in the SNc to DLS pathway^12,39–41^. Increased DA release in DLS, which is associated with general motor invigoration early in learning^8,42^, now increases the likelihood of initiating the previously learned, stereotyped motor plan^10,36^. Furthermore, inhibition of the DLS severely inhibits execution of a well-learned behavior^10^, while activation of DA release in DLS impairs reversal learning^60^. These data suggest that the MS’s reduction of SNc DA population activity may be improving strategy switching by decreasing the likelihood of initiating the action sequence associated with the well learned strategy. In support of this, MS activation specifically decreased perseverative errors, as well as increased choice latency. We also posit that this is the reason why MS activation was not effective until the initial strategy had been sufficiently well learned; i.e., the MS isn’t able to exert an influence over strategy switching until executing the strategy becomes dependent on the SNc to DLS pathway.

#### Are the DA population activity changes in VTA irrelevant to the MS’s ability to affect strategy switching?

Although intra-vSub infusion of scopolamine did not affect the MS’s improvement in strategy switching, it is important to note that infusion of both drugs inhibited the improvement in strategy switching to a greater degree than bicuculline alone. This suggests that scopolamine’s inhibition of the MS activation-induced increase in VTA DA population activity may have had some effect on the MS’s improvement in strategy switching, albeit a secondary effect. We posit that once a strategy has been sufficiently well-trained, altering DA activity in the SNc is the more impactful result of MS activation, as explained above. However, a secondary result of decreasing the likelihood of initiating the well-learned strategy/action sequence could be allowing a return of behavior to a “goal-directed” state. Once back in the goal-directed state, the MS’s increase in VTA DA activity may still provide some enhancement in learning rate for the new strategy^9^. Others have found that VTA DA release in VS increases early in reversal learning and is necessary to perform the reversal^43,44^, which supports the notion that increases in VTA DA activity/release could have been beneficial here.

## Conclusions

This study demonstrates that MS activation improves strategy switching once the initial strategy is sufficiently well learned. Interestingly, this effect is demonstrated as a slowing of decision-making, reducing perseverative errors, in order to adapt to a new strategy more quickly. This is primarily due to the MS’s reduction of DA activity in the SNc, a region that is responsible for initiating a well-learned action sequence^10,36^. The ability to flexibly overcome well learned or habitual thoughts and behaviors is impaired in a multitude of psychiatric disorders^45–54^. For example, addiction may stem from excessive habit formation leading to a loss of flexible control over behavior^54–57^, although this is contested by others^58,59^. This could lead to the inability of a person to cease drug-taking despite negative, often serious, consequences. The MS’s ability to downregulate SNc DA activity, as a means to destabilize the initiation of a well-learned action sequence and increase flexibility, could have the potential to be therapeutically-relevant in the study of this aspect of addiction. Thus, continued examination of the MS’s ability to promote flexibility will be important, and may have the potential to reveal novel treatment avenues in future studies.

