## Supplemental Figures for "Medial septum activation improves strategy switching once strategies are well learned via bidirectional regulation of dopamine neuron population activity"

**Supplementary Figures:**

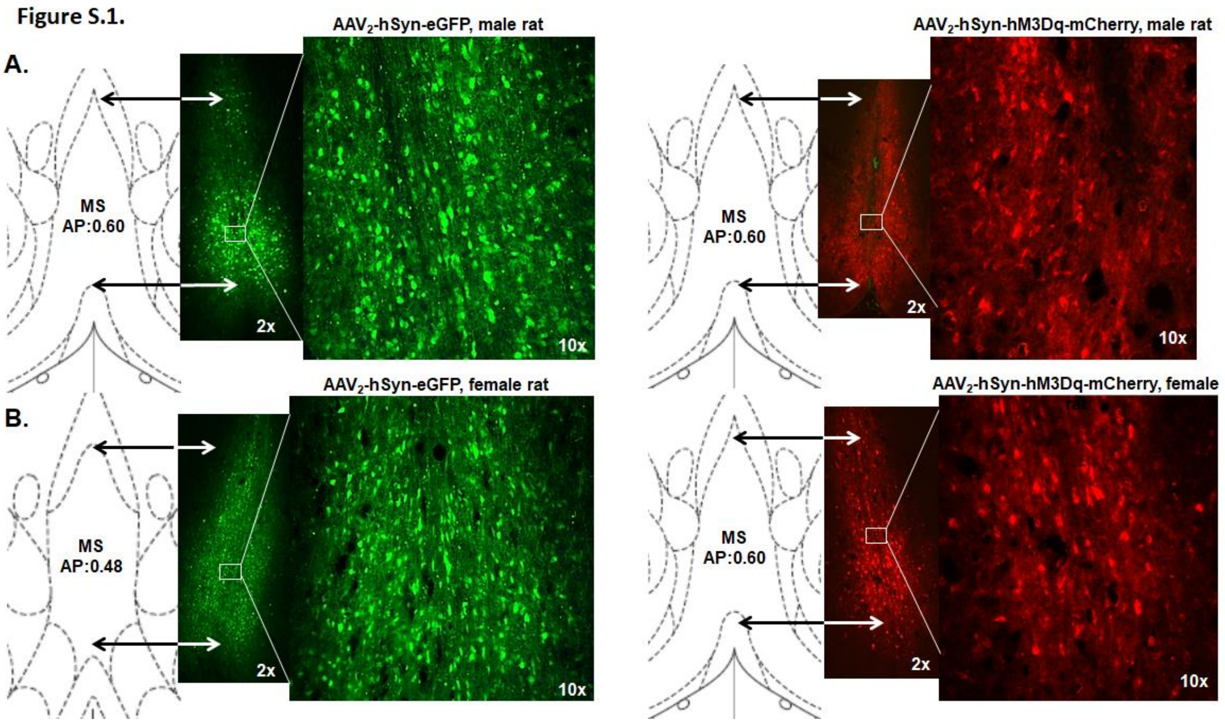


Figure S.1: Additional representative micrographs. Additional examples of **A.** male and **B**. female rats that were infused with an hM3Dq-containing (DR, AAV2 – hSyn – hM3Dq – mCherry) or an empty vector control (Con, AAV2-hSyn EGFP) virus into the medial septum (MS, AP: +0.6, ML: $\pm$0.5, DV: -6.1). Coronal section drawing indicates where the fluorescence signal was found within the MS (2x) and 10x shows expression in neurons within the MS.

**Figure S.2.**

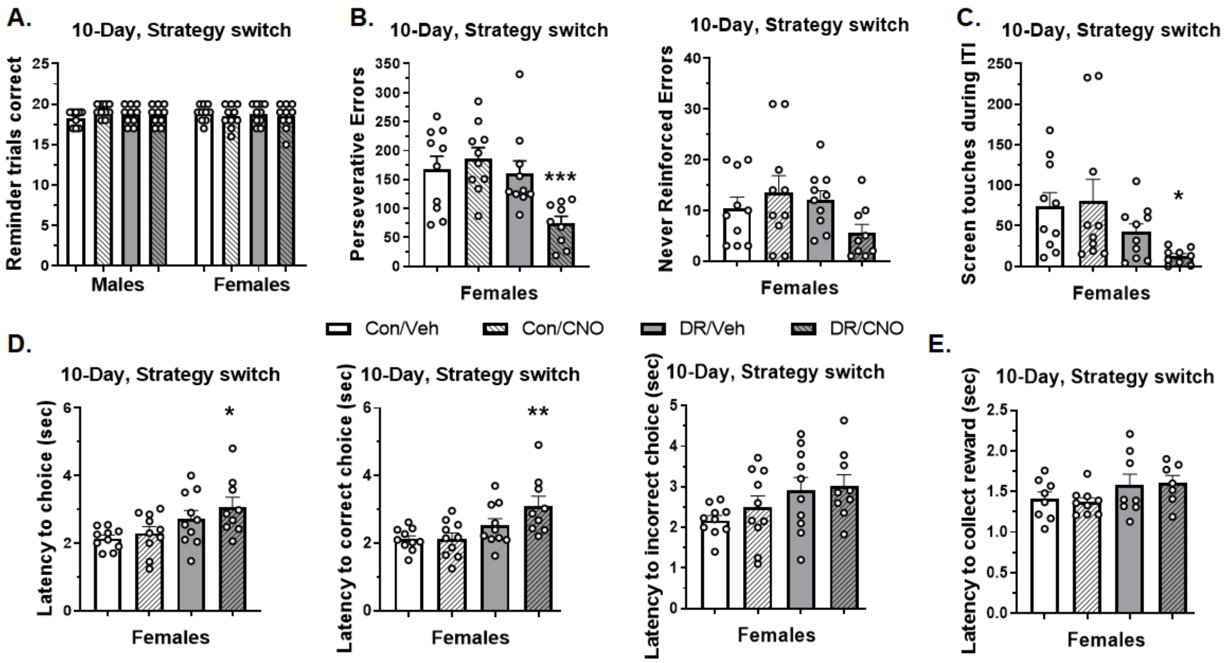


<u>Figure S.2: Reminder trials, error breakdown, and response latencies.</u> **A**. All rats performed similarly in the 20 "reminder" trials at the beginning of the session, suggesting that rats remembered the egocentric strategy to a similar degree (trials correct out of 20- males: Con/Veh: 18.2±0.3, Con/CNO: 19.2±0.2, DR/Veh: 18.7±0.4, DR/CNO: 18.7±0.4; females: Con/Veh: 18.9±0.3, Con/CNO: 18.5±0.4, DR/Veh: 18.8±0.4, DR/CNO: 18.5±0.5; $F_{7,71}$=0.60, P=0.75). **B.** Analysis of error type revealed that female DR/CNO rats committed fewer perseverative (Con/Veh: 167.8±22.4, Con/CNO: 186.7±18.6, DR/Veh: 160.1±22.1, DR/CNO: 74.1±12.4; $F_{3,35}$=6.16, P=0.002), but not never-reinforced (Con/Veh: 10.4±2.2, Con/CNO: 13.5±3.4, DR/Veh: 12.1±1.8, DR/CNO: 5.6±1.7; $F_{3,35}$=1.98, P=0.13), errors. ***A post-hoc Tukey's test revealed that perseverative errors were reduced compared to all three control groups (vs. Con/Veh P=0.011, vs. Con/CNO P=0.002, vs. DR/Veh P=0.021). **C.** Female rats also showed a significant reduction in impulsive screen nose-pokes during the inter-trial interval (Con/Veh: 73.6±17.3, Con/CNO: 80.0±27.3, DR/Veh: 42.1±10.3, DR/CNO: 12.4±3.2; $F_{3,35}$=3.07, P=0.041). *However, a post-hoc Tukey's test showed only a trend toward a reduction compared to the Con/Veh (P=0.089) and Con/CNO (P=0.051) groups. **D.** Female DR/CNO rats showed a significant increase in the time to make a correct (Con/Veh: 2.1±0.1, Con/CNO: 2.1±0.2, DR/Veh: 2.5±0.2, DR/CNO: 3.1±0.3; $F_{3,35}$=5.37, P=0.0038), but not incorrect (Con/Veh: 2.2±0.1, Con/CNO: 2.5±0.3, DR/Veh: 2.9±0.3, DR/CNO: 3.0±0.3; $F_{3,35}$=2.14, P=0.11), choice. **A post-hoc Tukey's test revealed that correct choice latency was increased compared to the Con/Veh (P=0.007) and Con/CNO (P=0.008) groups, but not the DR/Veh group (P=0.19). **E.** These changes occurred without affecting the latency to collect the pellet reward (Con/Veh: 1.4±0.1, Con/CNO: 1.4±0.1, DR/Veh: 1.6±0.1, DR/CNO: 1.6±0.1; $F_{3,28}$=1.48, P=0.24).

**Figure S.3.**

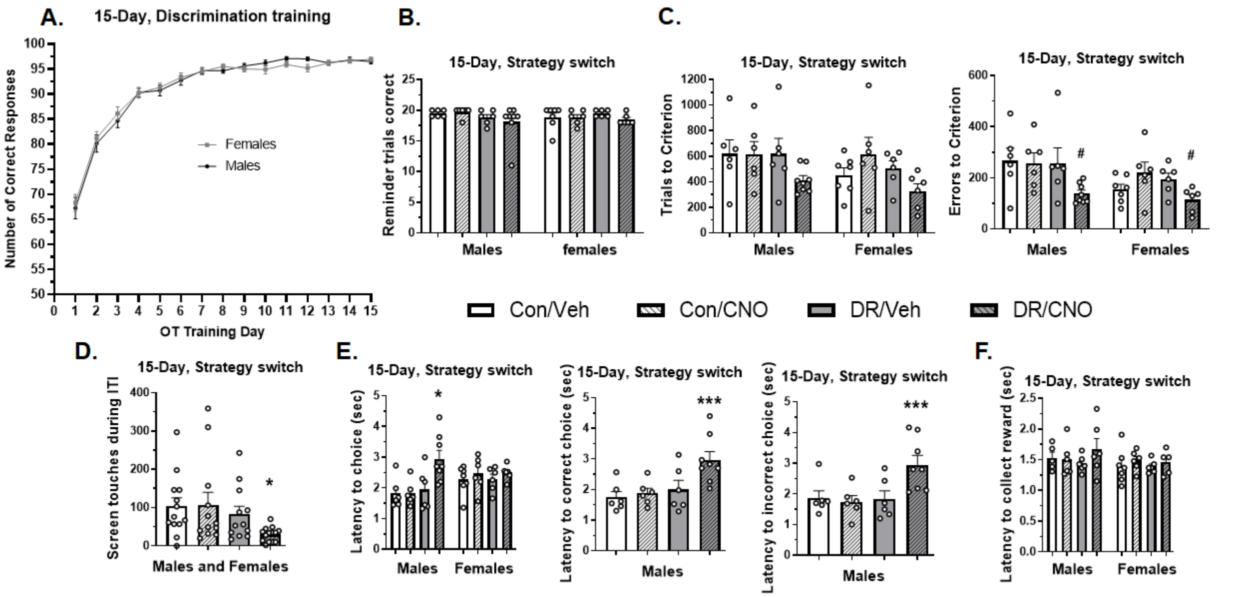


Figure S.3: Discrimination training, reminder trials, error breakdown, and response latencies. **A.** In contrast to the 10-day training rats, learning rates were not significantly different between sexes with 15 days of discrimination training. **B.** Rats again performed correctly on the substantial majority of the 20 “reminder” trials at the beginning of the session (trials correct out of 20- males: Con/Veh: 19.5±0.2, Con/CNO: 19.7±0.7, DR/Veh: 18.8±0.5, DR/CNO: 18.1±1.0, $F_{3,21}$=1.05, P=0.39; females: Con/Veh: 18.9±0.7, Con/CNO: 18.8±0.5, DR/Veh: 19.5±0.2, DR/CNO: 18.5±0.3, $F_{3,21}$=0.67, P=0.57). **C.** Trials to criterion and errors separated by sex. Chemogenetic activation of the MS reduced trials to criterion in males (N=6-8 rats/ group, Con/Veh: 618.8±109.9, Con/CNO: 615.5±100.1, DR/Veh: 618.0±121.6, DR/CNO: 413.1±35.5, $F_{3,22}$=1.44, P=0.26) and females (N=6-7 rats/ group, Con/Veh: 453.4±56.4, Con/CNO: 617.8±130.5, DR/Veh: 504.7±58.4, DR/CNO: 328.5±56.6, $F_{3,21}$=2.16, P=0.12), but neither reached statistical significance until combined. Chemogenetic activation of the MS reduced errors in males (Con/Veh: 266.8±49.7, Con/CNO: 257.8±40.5, DR/Veh: 257.7±59.6, DR/CNO: 138.6±13.6, $F_{3,22}$=2.53, P=0.083) and females (Con/Veh: 155.7±19.6, Con/CNO: 220.0±41.6, DR/Veh: 195.5±23.7, DR/CNO: 114.8±19.6, $F_{3,21}$=2.80, P=0.065), but both remained at #trend significance levels until combined. **D.** When combined, both sexes of the DR/CNO group showed a significant reduction in impulsive screen nose-pokes during the inter-trial interval (Con/Veh: 103.8±22.0, Con/CNO: 107.2±32.7, DR/Veh: 83.0±20.4, DR/CNO: 29.9±5.04, $F_{3,47}$=2.92, P=0.044). **E.** Choice latency was significantly increased in male, but not female, rats. However, this was seen when the rats made both correct (Con/Veh: 1.8±0.2, Con/CNO: 1.9±0.2, DR/Veh: 2.0±0.3, DR/CNO: 3.0±0.3; $F_{3,22}$=5.62, P=0.005) and incorrect (Con/Veh: 1.9±0.2, Con/CNO: 1.7±0.2, DR/Veh: 1.8±0.3, DR/CNO: 3.0±0.3; $F_{3,22}$=5.24, P=0.007) choices, suggesting a general slowing of decision making without it necessarily being beneficial. ***A post-hoc Tukey’s test revealed that choice times were reduced compared to all three control groups for both correct and incorrect choices, respectively (vs. Con/Veh P=0.009 and 0.034, vs. Con/CNO P=0.021 and 0.016, vs. DR/Veh P=0.046 and 0.028). **F.** Reward collection latency was unaffected by MS activation in either males (Con/Veh: 1.5±0.1, Con/CNO: 1.5±0.1, DR/Veh: 1.4±0.1, DR/CNO: 1.7±0.2; $F_{3,19}$=0.74, P=0.54) or females

(Con/Veh: 1.4$\pm$0.1, Con/CNO: 1.5$\pm$0.1, DR/Veh: 1.4$\pm$0.04, DR/CNO: 1.5$\pm$0.1; $F_{3,20}=0.38$, $P=0.77$).

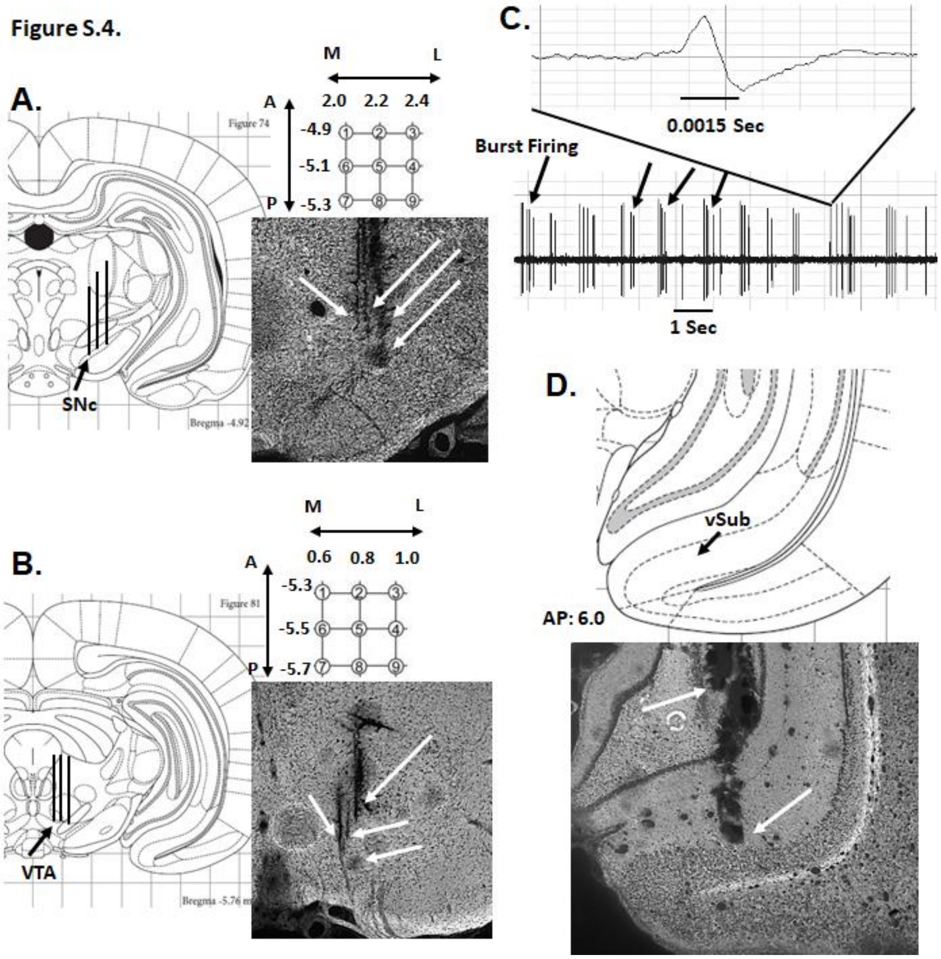


<u>Figure S.4: Representative recording electrode placement, recording, and vSub placement.</u> Recording microelectrodes constructed from borosilicate glass capillary tubes and filled with 2% Pontamine sky blue dissolved in 2.0M NaCl were lowered in 9 sequential tracks arranged in a predetermined grid pattern spaced at 0.2mm intervals in the **A.** substantia nigra pars compacta (SNc) and **B.** ventral tegmental area (VTA). Arrows depict actual electrode tracks in representative photomicrographs, as well as Pontamine sky blue placement mark. **C.** DA neurons were identified and recorded for 1-3 minutes. The total number of spontaneously active DA neurons within each animal was counted and then normalized by dividing by the total number of tracks that were examined (active DA neurons per electrode recording track or DA neurons/ track). **D.** Placement map showing the termination of the cannula location in vSub and corresponding representative photomicrograph. The top arrow marks the bottom of the guide cannula, and the bottom arrow marks the bottom of the infusion cannula. Rats with cannula placements outside the vSub were excluded.

Figure S.5.

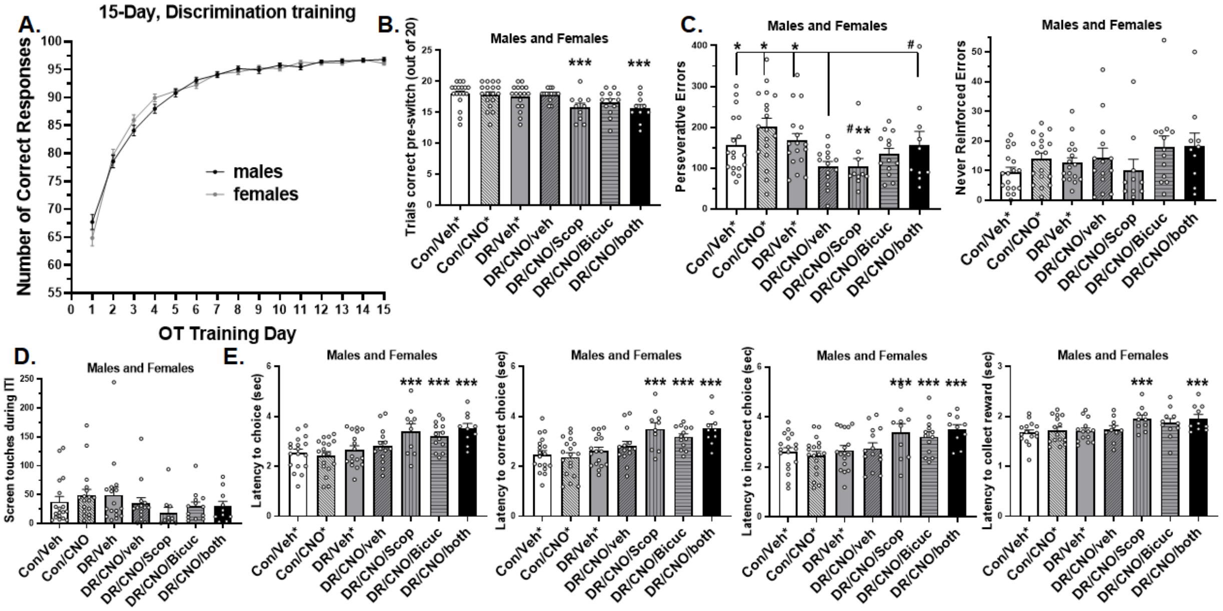


Figure S.5: Discrimination training, reminder trials, error breakdown, and response latencies. **A.** Similar to the previous experiment, learning rates were not significantly different between sexes with 15 days of discrimination training. **B.** All rats correctly performed the majority of the 20 “reminder” trials at the beginning of the session (trials correct out of 20: Con/Veh*: 17.9±0.5, Con/CNO*: 17.8±0.5, DR/Veh*: 17.5±0.3, DR/CNO/Veh: 17.9±0.3, DR/CNO/Scop: 15.8±0.7, DR/CNO/Bicuc: 16.6±0.6, DR/CNO/both: 15.6±0.7). Rats in the DR/CNO/Scop and DR/CNO/both groups showed a small but significant ($F_{6,93}$=3.316, P=0.005), reduction in number of correct trials out of 20. ***Pre-switch trials correct were reduced for both the DR/CNO/Scop and DR/CNO/Both groups, respectively (vs. Con/Veh* P=0.006 and 0.003, vs. Con/CNO* P=0.009 and 0.004, vs. DR/Veh* P=0.032 and 0.017, vs. DR/CNO/Veh P=0.012 and 0.006). **C.** The reduction in perseverative errors after MS activation was prevented by infusion of both bicuculline and scopolamine (Con/Veh*: 156.3±17.1, Con/CNO*: 201.3±20.9, DR/Veh*: 167.1±17.6, DR/CNO/Veh: 104.0±12.2, DR/CNO/Scop: 104.3±19.2, DR/CNO/Bicuc: 134.2±14.4, DR/CNO/both: 156.8±33.1; $F_{6,93}$=3.33, P=0.005). Infusion of bicuculline or scopolamine alone did not affect the reduction in perseverative errors (vs. DR/CNO/Veh P=0.28 and 0.99, respectively). *#A Fisher’s LSD test showed that the DR/CNO/Veh and DR/CNO/Scop groups were reduced compared to the Con/Veh* (P=0.048 and 0.075, respectively), Con/CNO* (P=0.0003 and 0.001, respectively), DR/Veh* (P=0.019 and 0.034, respectively), and DR/CNO/Both (P=0.085 and 0.11, respectively). Never-reinforced errors were not affected by any systemic or intra-vSub treatment (Con/Veh*: 9.5±1.6, Con/CNO*: 14.1±1.8, DR/Veh*: 12.6±1.7, DR/CNO/Veh: 14.3±3.2, DR/CNO/Scop: 10.0±3.8, DR/CNO/Bicuc: 18.0±3.7, DR/CNO/both: 18.3±4.3, $F_{6,93}$=1.51, P=0.18). **D.** Nose pokes during the intertrial interval (ITI) were not affected by any systemic or intra-vSub treatment (Con/Veh*: 37.6±9.3, Con/CNO*: 49.0±10.1, DR/Veh*: 49.2±13.9, DR/CNO/Veh: 34.7±10.0, DR/CNO/Scop: 18.8±9.1, DR/CNO/Bicuc: 30.1±7.2, DR/CNO/both: 29.8±8.6, $F_{6,93}$=1.00, P=0.43). **E.** Infusion of scopolamine, bicuculline, or both in the DR/CNO, but not control, rats led to a slight, but statistically significant, increase in choice (Con/Veh*: 2.5±0.2, Con/CNO*: 2.4±0.2, DR/Veh*: 2.7±0.2, DR/CNO: 2.8±0.2, DR/CNO/Scop: 3.4±0.3,

DR/CNO/Bicuc: 3.2±0.2, DR/CNO/both: 3.5±0.2; $F_{6,93}=5.32$, $P<0.0001$) and reward collection times (Con/Veh*: 1.7±0.1, Con/CNO*: 1.7±0.1, DR/Veh*: 1.7±0.1, DR/CNO: 1.7±0.1, DR/CNO/Scop: 2.0±0.1, DR/CNO/Bicuc: 1.9±0.1, DR/CNO/both: 2.0±0.1; $F_{6,78}=2.71$, $P=0.019$) compared to the other 4 groups (***Fisher's LSD all p's $< 0.05$).
